# Fezf2 and Aire1 evolutionary trade-off in negative selection of T cells in the thymus

**DOI:** 10.1101/2022.02.01.478624

**Authors:** Michel-Edwar Mickael, Norwin Kubick, Agata Gurba, Pavel Klimovich, Irmina Bieńkowska, Tomasz Kocki, Mariusz Sacharczuk

## Abstract

In vertebrates, thymus expression of various body proteins to eliminate autoreactive T cells during the negative selection process is orchestrated by AIRE and FEZF2. T cells first appeared in vertebrates. However, the evolutionary history of these two genes in relation to T cells emergence is still not clear. Specifically, it is still not known, whether these two genes emerged concurrently to support the negative selection process. Furthermore, whether there is an evolutionary trade-off between these two genes is not known. Whether these two genes play a similar role in controlling auto-reactivity elimination in lampreys and invertebrates is also unknown. We used a plethora of phylogenetic analysis tools including; multiple sequence alignment, phylogenetic tree building, ancestral sequence reconstruction, functional specificity investigation, and positive selection analysis to address these questions. We found that these two genes represent two distinct pathways of negative selection with two unique origins. While AIRE emerged during the divergence of T cells in vertebrates, FEZF2 is far ancient with homologs in invertebrates including Cnidaria, Trichoplax. We found that FEZF2 structure is highly conserved between invertebrates and vertebrates. Moreover, the genes controlled by both families included a mixture of ancient and recently diverged genes. However, we found that AIRE contains an LXXLL motif for binding nuclear receptors. Conversely, FEZF2 possesses several motifs known to play a role in autophagy, such as DKFPHP, SYSELWKSSL, and SYSEL. However, both genes contain similar motifs such as MAPK regulating motifs. Interestingly, AIRE seems to be lacking in lampreys, in contrast to FEZF2. Taken together, our investigation hints that FEZF2 was initially employed to control a rudimentary auto-reactivity elimination process in invertebrates, then evolved to play a part in controlling a negative selection process in early vertebrates and higher vertebrates. The emergence of AIRE seems to be correlated with controlling the negative selection process in higher vertebrates. The results demonstrate a strong evolutionary trading-off process, where FEZF2 kept controlling certain biological processes whereas AIRE gained control of others. Several critical genes are controlled by both genes, to ensure an adequate negative selection process.

## Introduction

The negative selection process aims to eliminate autoreactive T cells that attack the body’s own cells (1) (2). During T cells development in the thymus, each CD4+CD8+ αβ T cell express a TCR generated by V(D)J recombination (3). Positive selection process occurs when CD4+CD8+T cells acquire survival signals to differentiate into single-positive T cells according to their MHC-TCR affinity (4). Subsequently, single positive T cells are recruited to the medulla. This migration process is controlled by the interaction between CCR7 expressed on T cells and its ligands (e.g., CCL19 and CCL21) expressed by medullary thymic epithelial cells (mTECs) (5). Negative selection occurs in the medulla where CD4 or CD8 T cells interact with antigen-presenting cells, such as mTECs and dendritic cells (6). mTECs express various tissue-restricted antigens (TRAs), to allow deletions of T cells specific for antigens that otherwise would only be encountered in the periphery. Almost all T cells recognizing self-antigen-MHC complexes are deleted by mTECs and thymic dendritic cells (3). T cells expressing a functional TCR without significant reactivity to self-antigens migrate to secondary lymphoid organs (e.g., spleen and lymph nodes) and circulate throughout the body.

The negative selection process is controlled by two main proteins, namely AIRE1 and FEZF2 (3). In the mouse thymus, AIRE1 controls 40 % of TRA expression (7) (8). AIRE1 does not seem to be a transcription factor(3). Rather, AIRE1 induces the transcription of TRA genes through interactions with various transcriptional factors. These interactions include enrichment of repressive markers in its promoter region (e.g., H3K27) (3). It also supports transcriptional elongation through facilitating p-TEFb recruitment. Additionally, AIRE1 interacts with BRD4 which is a transcriptional and epigenetic regulator(9). AIRE1 also interacts with TOP1 and TOP2 which are known for their role in regulating the topologic states of DNA during transcription (10). AIRE1 expression is controlled by various gene networks, such as the TNFR family network. This network includes RANK, CD40, RANK-ligand, and CD40-ligand as well as NF-κB pathways (11). It was also shown that estrogen and androgen perform a crucial role in controlling AIRE1 expression (12). Recently, strong evidence emerged supporting a crucial role of FEZF2 in central tolerance, probably through acting as a transcription factor. A difference in the usage of TCR Vβ chains in CD4 or CD8 T cells between WT and FEZF2-deficient thymus demonstrated that FEZF2 regulates the negative selection of T cells (13). Notably, the genes controlled by FEZF2 are not controlled by AIRE1. However, some exceptions exist such as Fabp2, Saa2, and Cdkn1c. FEZF2 expression itself is controlled by LTBR signaling pathway. If other genes are fundamental for the process of central selection is not currently known (3).

The evolutionary history of FEZF2 and AIRE1 is still unknown. FEZF2 belongs to the FEZF family. This family functions as a transcriptional repressor, and it is known to contain six C2H2 zinc fingers and an EH1 repressor motif (14)(15). FEZF2 function is not limited to the crucial process of negative selection in the thymus, it is also expressed in the brain, where it plays a role in neural development by repressing HES5 (14). However, whether it plays other roles is unknown. How FEZF2 evolved in relation to T cells emergence is still not clear. AIRE1 contains four crucial motifs that support its function in controlling the function of various TFs(3). These motifs are namely CARD (caspase recruitment domain), SAND (Sp100, Nucp41/p75, and Deaf1), PHD1 (plant homeodomain 1), and PHD2. Although AIRE1 is known to exist in vertebrates, whether it has any homologs in invertebrates is still unknown. Whether AIRE co-evolved with T cells to support the negative selection process is still unclear. The evolutionary advantage of having more than one gene controlling the process of negative selection has not been yet been identified. Also, the evolutionary origin of the negative selection process remains elusive. Whether this process specifically evolved to eliminate autoreactive T cells or whether it mirrors simpler models in invertebrates is not clear.

In this research, we focused on studying the evolutionary history of the currently known genes that control the negative selection process. In order to achieve this task, we performed an array of phylogenetic analyses, including multiple sequence alignment, followed by tree building. Subsequently, we analyzed the origins of FEZF2 and AIRE1 through using ancestral sequence construction. we also investigated their positive selection using PAML. In order to compare functional specifications, we used DIVERGE and linear functional motifs search, as well as gene enrichment analysis for their downstream targets. The results of our research permitted us to infer the origin and the evolution of the negative selection mechanism.

### Database search

We focused in this research on investigating the FEZF2 and AIRE1 Pathways and their evolutionary origins. To draw a full picture of these two pathways, we investigated FEZF2, AIRE1 as well as their regulators and identified downstream targets (3). Due to the diverse nature and long evolutionary history of the investigated genes we employed protein sequence alignment (16). Moreover, to make sure that our analysis is a reasonable representation of the dynamic evolutionary process of these two genes history, we examined the presence of FEZF2 and AIRE1 in rodents, Actinopterygii, Arthropods, Spiralia, Tunicates, Cnidaria and Placozoa, spanning more than 500 million years. Human protein families were utilized for BLASTP searches against the aforementioned proteomes. The longest transcript homolog was employed in the investigation. Resulting sequences were accepted as candidate proteins if their E values were lower than 1e^-10^(17). Sequences were additionally filtered by comparing the conserved domain in each protein against the human proteins investigated.

### Alignment and phylogenetic analysis

The phylogenetic investigation was done in two stages (18)(19). First, each investigated amino acid sequence was aligned using MAFFT through utilizing the iterative refinement method (FFT-NS-i)(20). Afterward that, we used, PHYML implemented in Seaview with five random starting trees to produce the final tree. The Neutrality test was accomplished using MEGA6 (21).

### Positive selection analysis

We utilized a maximum likelihood approach to investigate whether FEZF2 or AIRE2 experienced positive selection during evolution (22)(23). In the first instance, respective complementary DNAs (cDNAs) were back translated using EMBOSS Backtranseq tool and aligned according to their codon arrangement(24). Subsequently, we inspected positive selection patterns in both genes using CODEML (PAML)(25). To investigate selection we calculated the substitution rate ratio (ω)as the ratio between nonsynonymous (dN) to synonymous (dS) mutations. We utilized three different levels of investigation (i) Basic(global) selection (ii) branch,(iii) branch-site, and (iv) sites models(18). Statistical significance was calculated using a likelihood ratio test (LRT) based on the following equation; *P* value = χ^2^ (2*Δ(ln(LRT_model_) - ln(LRT_neutral_)), number of degrees of freedom).

### Linear motifs search

To explore the difference between FEZF2 and AIRE1 evolution-function association, we searched the protein sequences of both proteins for linear motifs. Linear motifs are defined as short sequences of amino acids that function as putative protein interaction sites. We implemented the search using the ELM server http://elm.eu.org/ with a motif cut-off value of 100 (26).

### Functional divergence estimation

Type II functional divergence with FEZF2 and AIRE1 proteins was investigated. The aim of this analysis to detect any shift of cluster-specific amino acid property. This type of divergence constitutes a shift of amino acid property (charge, size, hydropathy)(27). AIRE1 was clustered into higher and lower vertebrates, whereas FEZF2 was clustered in vertebrates and invertebrates (27)(18).

### Functional ontologies

We compared the gene enrichment of AIRE1 and FEZF2 downstream targets. This was done using three methods. First we analyzed the microarray gene sets GSE69105 and GSE2585 using GeneSpring©. These two public datasets contains group of WT and Knock out mice for AIRE1 and FEZF2. We accepted downstream genes candidates under the constraint of having a fold change >1.5, and p < 0.05. After that, we ran the gene ontology Gorilla server to identify gene enrichment in both datasets. In the second method, we used the gene list of downstream targets identified in (3) as an input to Metascape©. In the third method the genes identified in(3) were manually mined using GeneCards.

## Results

### Workflow

To investigate the evolutionary history of the proteins responsible for the negative selection pathway, we divided these proteins into three groups (i) main regulators (AIRE and FEZF2) (ii) upstream of the main regulators (e.g., regulators of regulators) (iii) downstream targets of the regulatory elements (Table 1). For the first group, we drew the phylogenetic history, calculated the ancestral sequence, investigated positive selection and functional divergence, functional specificity, and identified functional motifs. For the other two groups, we only investigated the evolutionary history and gene ontology (figure 1).

**Table 1.**
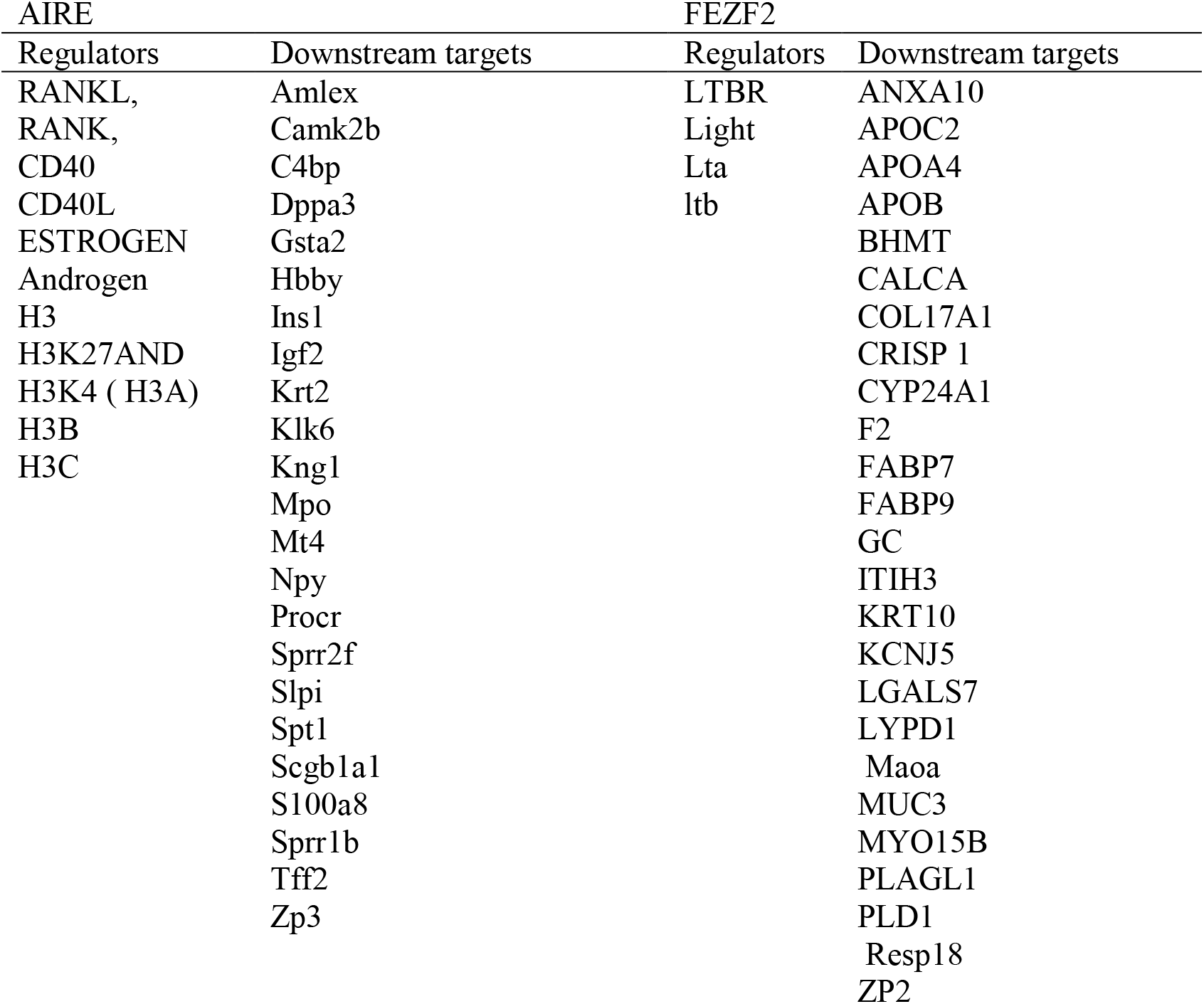
Investigated Genes classified based on their function in the selection process.

**Figure 1.**
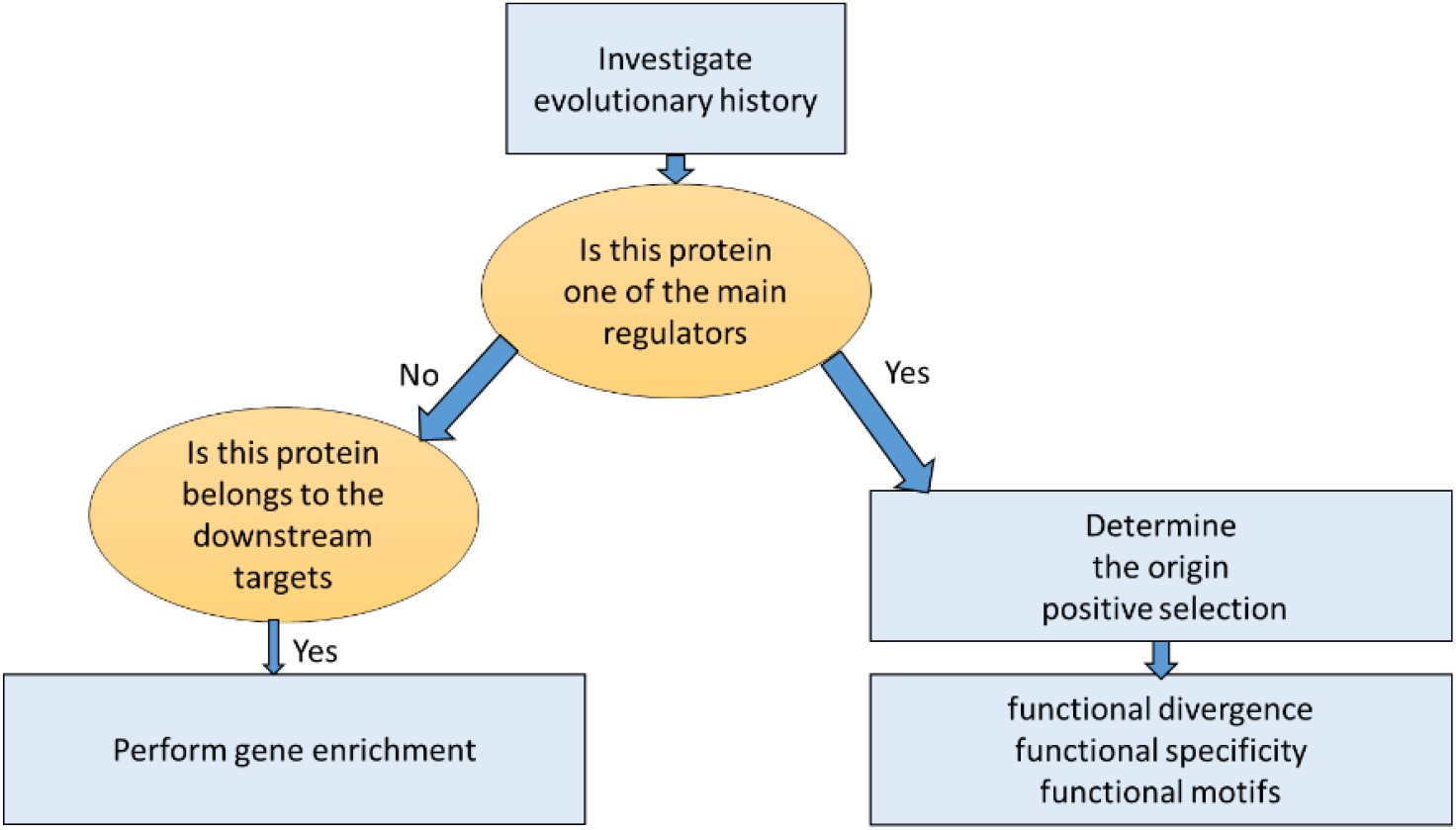
Workflow of the analysis. All proteins identified were phylogenetically investigated. The main regulators (e.g., AIRE1 and FEZF2) were investigated further by building their ancestral sequences, investigating their selection, divergence, specificity, and motifs. Downstream targets were examined for their significant gene enrichment pathways.

### 3.2. Evolutionary History of negative selection pathway molecules

The evolutionary history of the negative selection pathway proteins is complex. AIRE1 first emerged during bony fish divergence (figure 2). Interestingly, AIRE regulators do not share a specific emergence period. RANK first emerged during lampreys divergence. CD40 has homologs in both lampreys and Tunicate. CD40L seems to have converged later as it fit appeared in fish. Estrogen receptors (ER1 and ER2) both appear in lampreys. However, ER1 seems to be more ancient with a homolog in Spiralia and Cnidaria. Conversely, androgen seems to have emerged during bony fish divergence. Downstream targets of AIRE1 also do not share a similar evolutionary history. Various genes controlled by AIRE1 expression emerged in bony fish (e.g., CAMK2B, C4BP, KRT2) or rodents (e.g., AMPLEX). Similarly, various genes controlled by AIRE were as ancient as Cnidaria such as GSTA2, IGF2. The evolution of the FEZF2 pathways seems to follow that of AIRE, with no apparent common time of emergence. However, in contrast to AIRE, FEZF2 seems to possess ancient homologs in Trichoplax, Cnidaria, and Spiralia. Furthermore, several genes regulated by FEZF2 appeared in invertebrates. For example, BHMT has homologs in both Cnidaria and Trichoplax. F2 and MAOA emerged in Cnidaria while ITIH3 and KCNJ5 first appeared in Spiralia. Paradoxically current known regulators of FEZF2 seem to have diverged much later with LTA, LTB, and light diverging in bony fish, while LTBR first appearing in lampreys.

**Figure 2.**
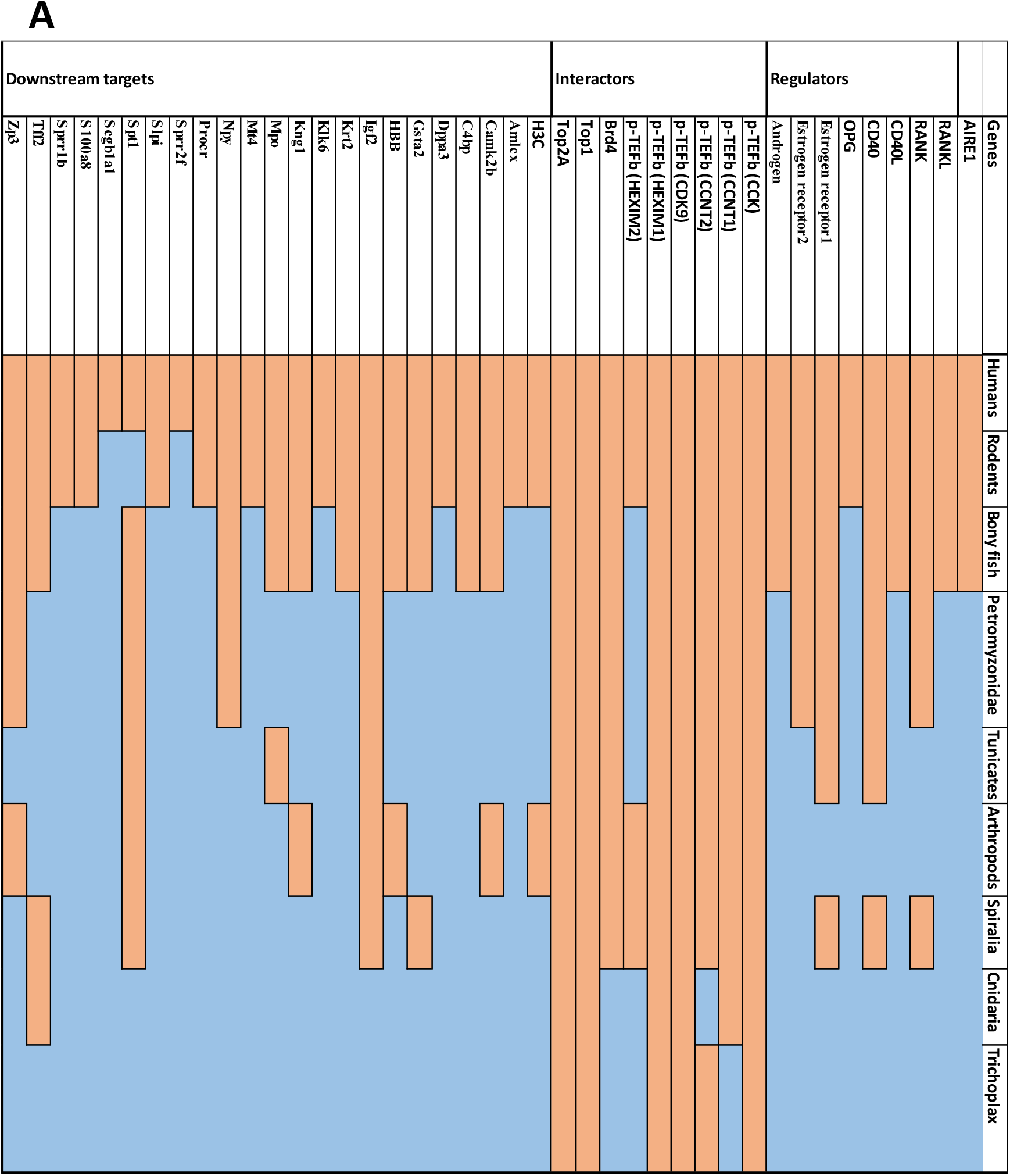
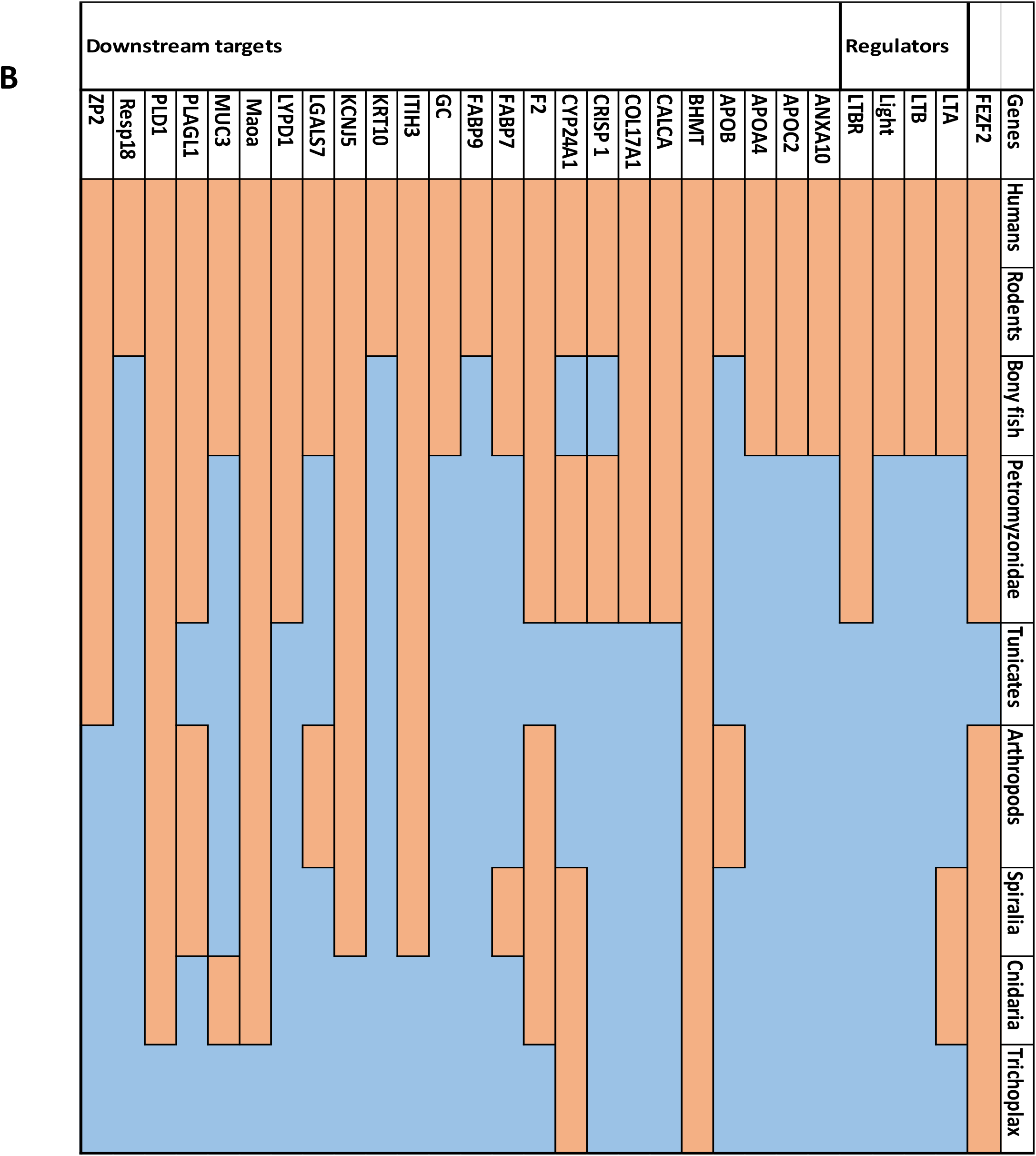
Evolutionary history of the AIRE and FEZF2 negative selection pathways. Our results indicate proteins involved in FEZF2 nor AIRE pathways do not seem to have emerged during a common evolutionary origin period. However, FEZF2 is more ancient than AIRE.

### Origin of regulatory proteins controlling the negative selection process

Regulatory elements controlling the process of negative selection have divergent evolutionary history (Table 2). The FEZ family contain three main genes, namely FEZ1, FEZ2 and UNC76. FEZF family seems to have diverged from a Zinc finger domain-containing protein, C2H2 type (*Metschnikowia aff. Pulcherrima*) (figure 3) (Table2). AIRE1 seems to have emerged from a PHD containing protein (figure 3).

**Table 2.**
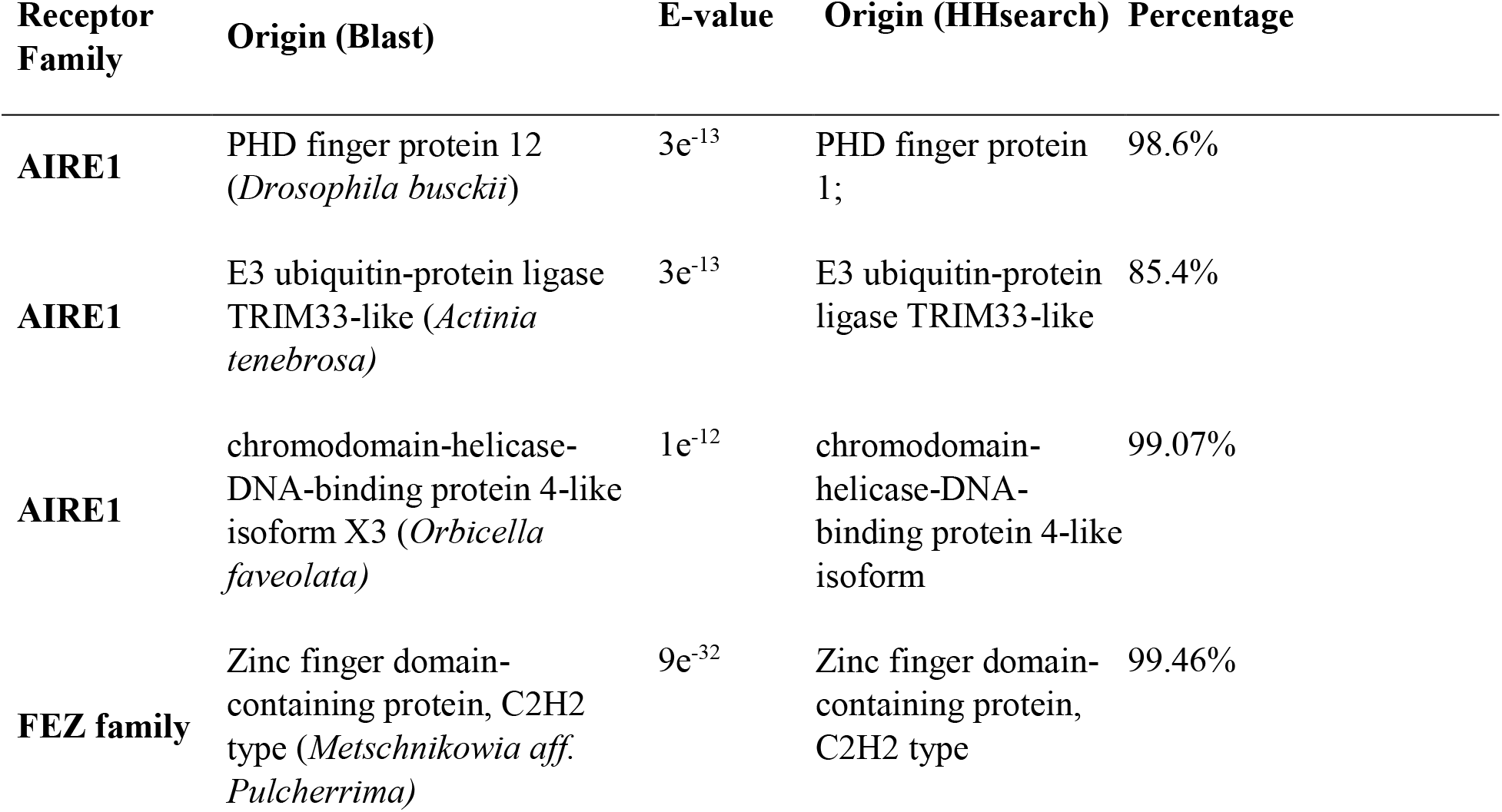
Estimated origins of AIRE complex compared to that of FEZF2.

| Receptor Family | Origin (Blast) | E-value | Origin (HHsearch) | Percentage |
| --- | --- | --- | --- | --- |
| AIRE1 | PHD finger protein 12 (*Drosophila busckii*) | $3e^{-13}$ | PHD finger protein 1; | 98.6% |
| AIRE1 | E3 ubiquitin-protein ligase TRIM33-like (*Actinia tenebrosa*) | $3e^{-13}$ | E3 ubiquitin-protein ligase TRIM33-like | 85.4% |
| AIRE1 | chromodomain-helicase-DNA-binding protein 4-like isoform X3 (*Orbicella faveolata*) | $1e^{-12}$ | chromodomain-helicase-DNA-binding protein 4-like isoform | 99.07% |
| FEZ family | Zinc finger domain-containing protein, C2H2 type (*Metschnikowia aff. Pulcherrima*) | $9e^{-32}$ | Zinc finger domain-containing protein, C2H2 type | 99.46% |

**Figure 3.**
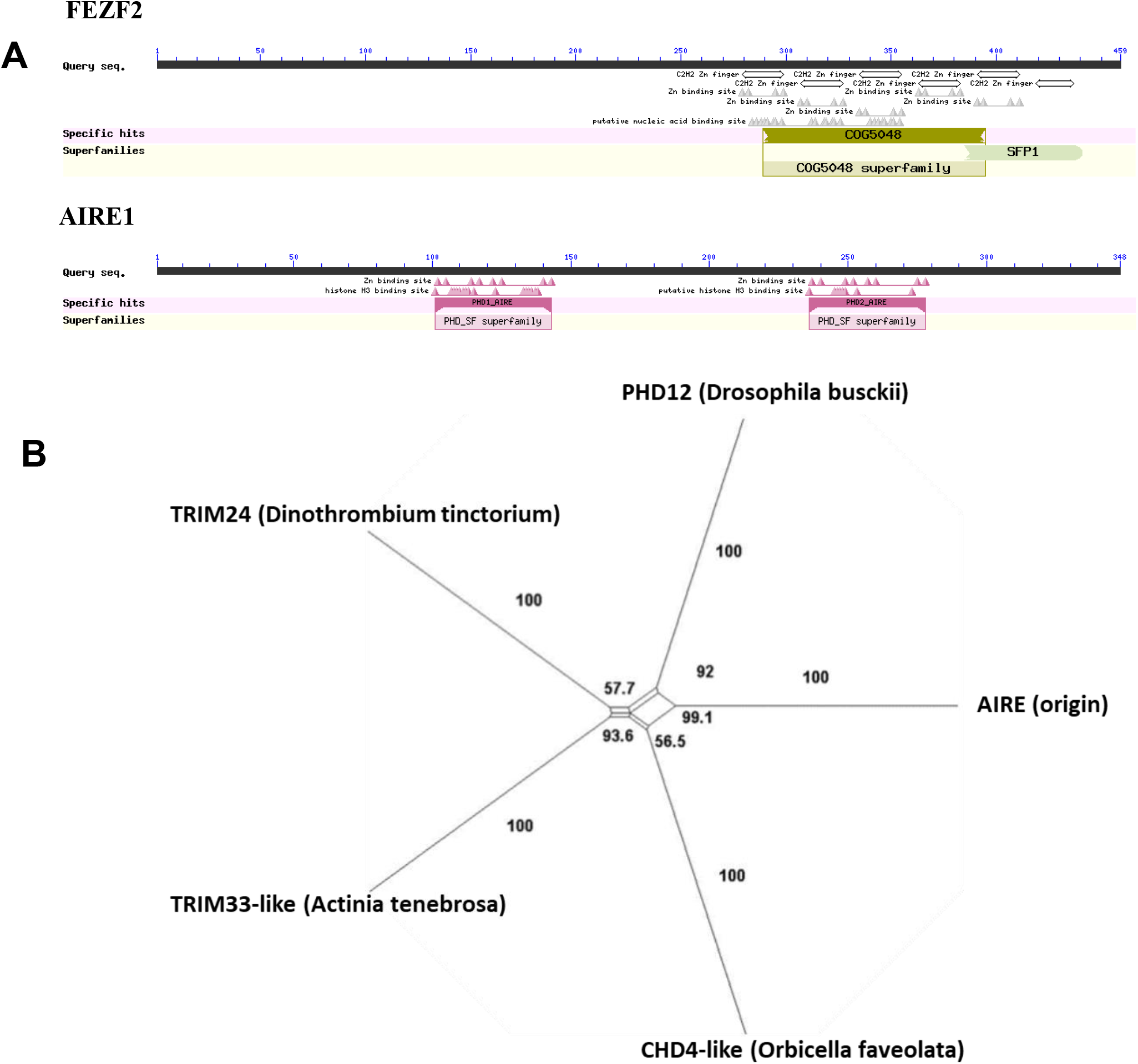
AIRE1 origin resolved using evolutionary network analysis. A) Structural analysis of the AIRE1 indicates that it possesses a PHD motif, while FEZF2 contains a CH2 Zinc finger motif. B) Our analysis indicates that AIRE1’s nearest homolog is PHD12 that first diverged in *Drosophila busckii*.

### Functional divergence

Overall a low degree of functional divergence was detected. We employed DIVERGE 3.0 to identify putative functional divergence type II sites using a cutoff value of 0.5. We detected only two functional divergent sites in AIRE1 sequence particularly at the 290 (where higher vertebrates have Histidine while bony fish have Tyrosine). Also, at amino acids (320,321), higher vertebrates have HA, while fish YS. Although the FEZ2 family has diverged much earlier than AIRE, we were able to locate only one single site that could represent a candidate for functional divergence (e.g., site 454) (figure 4). There is a strong variation at this site, where, vertebrates have Glutamine while insects have Threonine, Cnidaria have isoleucine, and Trichoplax has valine. These results suggest that both AIRE1 and FEZ2 express a limited number of functional divergent sites.

**Figure 4.**
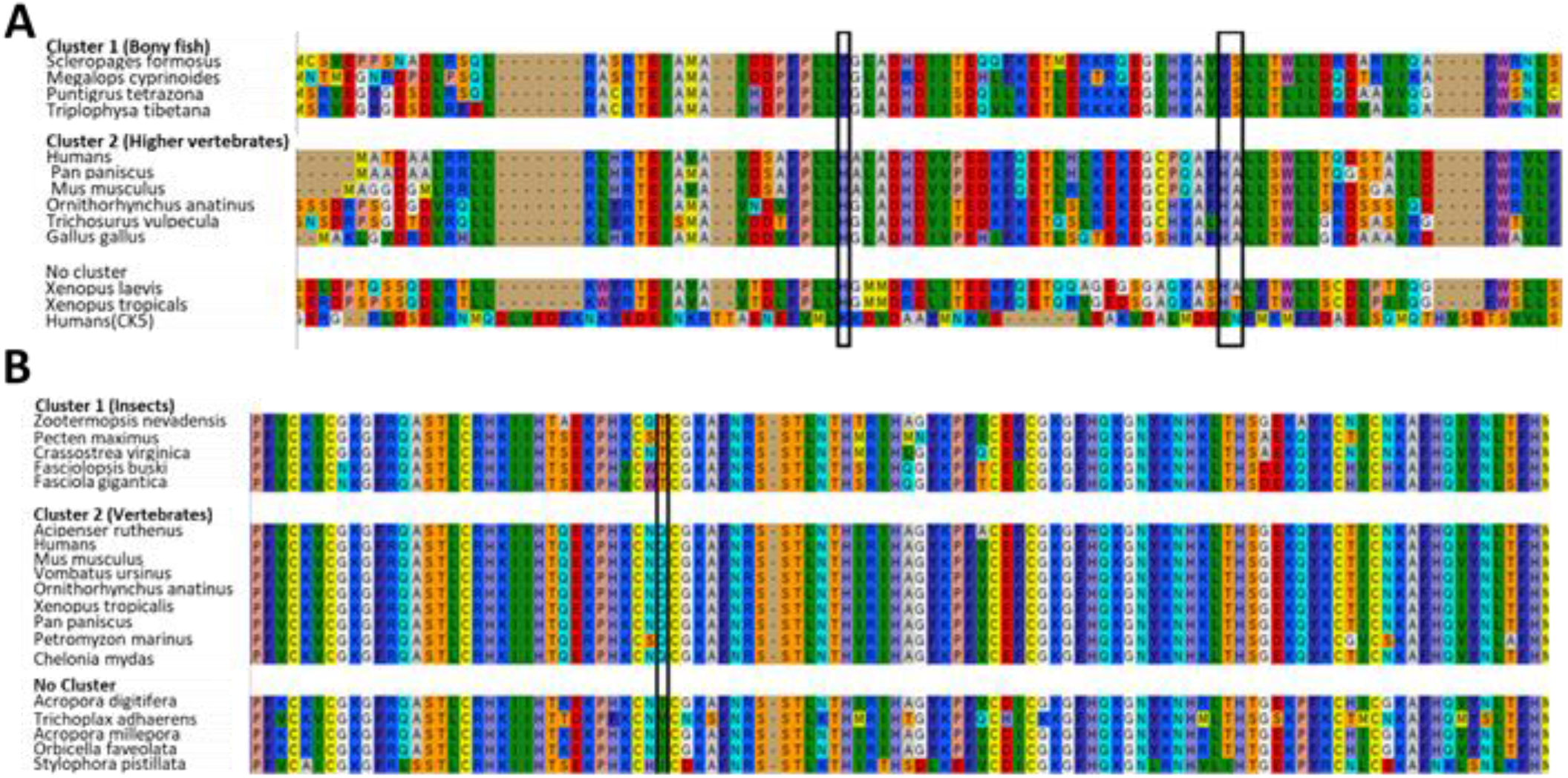
Functional divergence estimate for a) AIRE and b) FEZF2. Our analysis indicate that AIRE and FEZF2 possess a limited number of functional divergent sites based on type II divergence.

### 3.6. Neutrality Test and Positive Selection

We performed an analysis of positive selection for AIRE1 and FEZF2. Interestingly, we found that FEZF2 is highly conserved between vertebrates and invertebrates. This has been mirrored in an ω less than 1 (0.3, p-value<0.01) for both the global evolution pattern and the branch evolution pattern. Additionally, we did not identify any amino acids that were subjected to positive evolution on the branch-site level or the site levels based on the vertebrates and in-vertebrates divide (Table 3). In the case of AIRE1, we found that its conservation evolutionary pattern is more relaxed than FEZF2. Globally, AIRE1 seems to have evolved under neutral selection (ω=1). Similarly, we could not detect evidence of selection on the primates branch (Table 3). On the site branch, we found only two amino acids that were subjected to positive selection namely, 147 P (probability: 0.998**) and 315 V (probability: 0.992**). We performed neutrality tests for AIRE1 and FEZF2 by computing the D value (Tajima’s Neutrality Test). We found that both AIRE and FEZF2 show negative values indicating purifying conserved selection (Table 4).

**Table 3.** Positive selection test for the AIRE and the FEZ family.

| **Investigation** | **AIRE** | **FEZ** |
| --- | --- | --- |
| **Global** | 1.09 (p-value> 0.05) | 0.30830 (p-value <0.01) |
| **Branch** | 1.13 (p-value> 0.05) | 0.30852 (p-value <0.01) (done with lampreys, |
| **Site branch** | 147 P (probability : 0.998**)<br>315 V (probability: 0.992**)<br>(p-value> 0.05) | None |
| **Site** | Neu position<br>97 P 0.978*<br>129 A 0.996**<br>161 G 0.996**<br>170 R 0.995**<br>171 P 0.984*<br>178 A 0.950*<br>195 R 0.994**<br>215 E 0.957*<br>223 P 0.987* | None |

**Table 4.** Neutrality analysis comparing AIRE1 and FEZF2.

|  | **m** | **S** | **Ps** | **$\theta$** | **$\pi$** | **D** |
| --- | --- | --- | --- | --- | --- | --- |
| **AIRE1** | 10 | 1581 | 0.70 | 0.24 | 0.19 | -1.12 |
| **FEZF2** | 19 | 1283 | 0.53 | 0.15 | 0.11 | -1.07 |

$m$ = number of sequences, $n$ = total number of sites, $S$ = Number of segregating sites, $p_s = S/n$ , $\theta = p_s/a_1$ , $\pi$ = nucleotide diversity, and $D$ is the Tajima test statistic.

### Motifs search

Our linear motif search painted a complex picture for AIRE1 and FEFZ2. We found that AIRE1 contains an LXXLL motif which is known to play a role in binding nuclear receptors. Conversely, FEZF2 has several motifs that play a role in autophagy such as DKFPHP, SYSELWKSSL, and SYSEL, as well as actin and cytoskeleton dynamics regulation such as PPACPR. However, both genes contain various similar motifs such as MAPK regulating motifs (e.g., AGASPAT) (supplementary file 1).

### AIRE and FEZ have distinctive functional ontologies

We used the top genes detected by the microarray data (GSE69105 and GSE2585) analyzed in GeneSpring, with fold change >1.5, and p < 0.05) and used Gorilla server to predict gene enrichment pathways,(supplementary file 2) in addition to Metascape analysis (figure 5) and GeneCards mining (Table 5). Our Results indicate that FEZF2 and AIRE1 control multiple genes in a complementary fashion.

**Figure 5.**
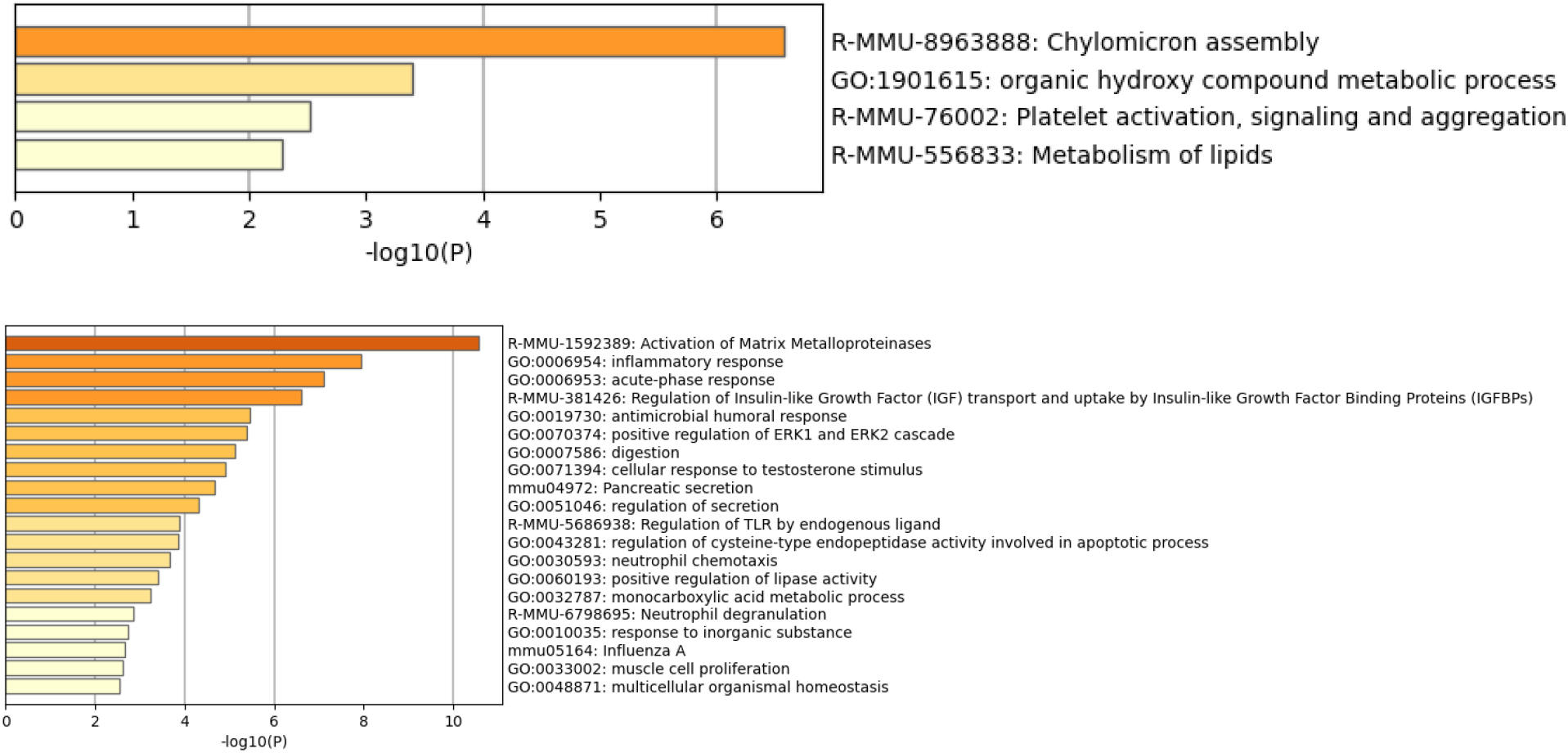
Metascape analysis of the downstream targets. Our analysis indicate that FEZF2 and AIRE sharing the control of the expression of various TRA pathways.

**Table 5.**
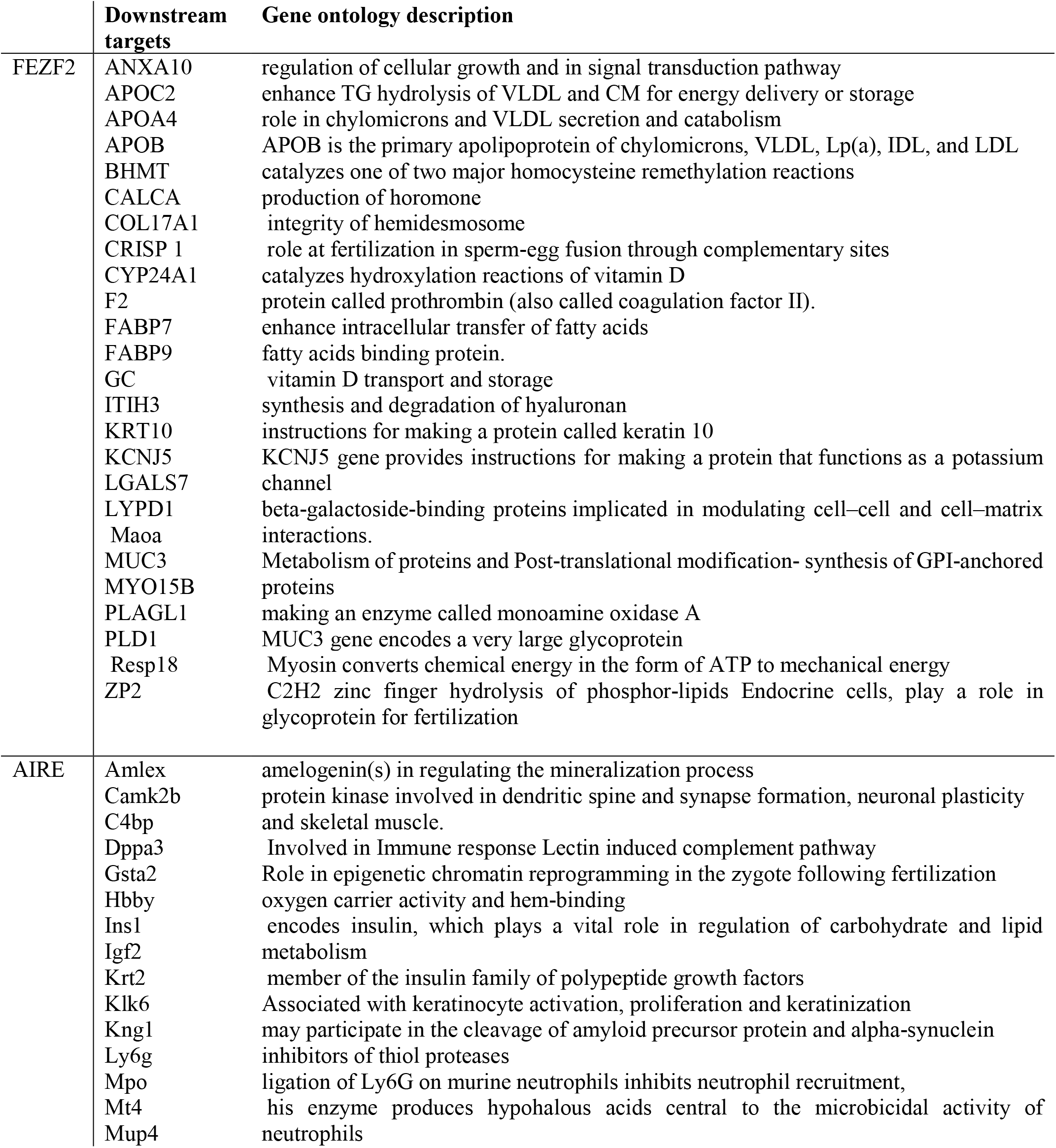
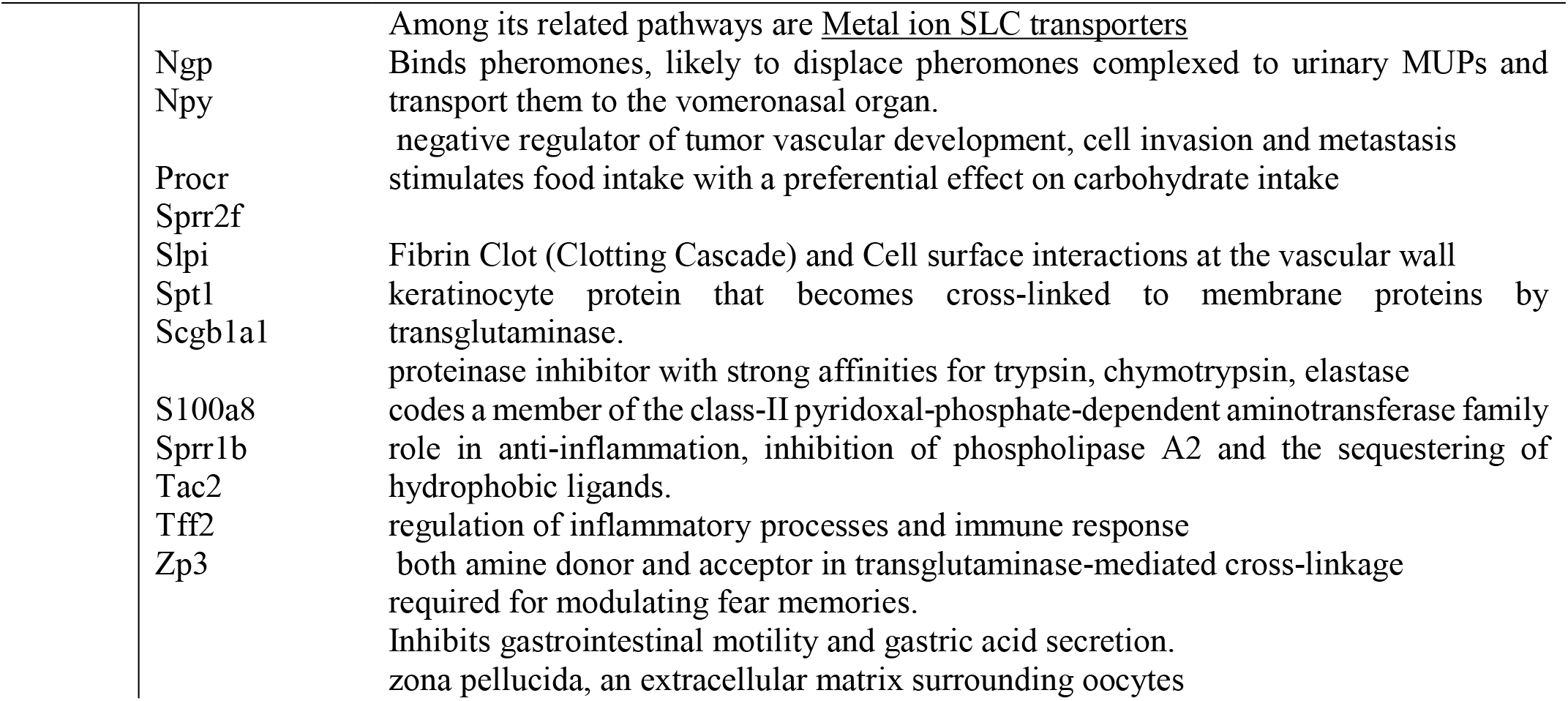
Gene ontology of the FEZF2 and AIRE controlled genes.

## 4. Discussion

Our analysis draws a complex analysis for the evolution of the negative selection of T cells in the thymus. Currently, two known genes are known to control the expression of various proteins that are needed to test T cell autoreactivity for eliminations. These two genes are AIRE and FEZF2. Our analysis shows that the AIRE pathway seems to be novel, while FEZF2 is more ancient with FEZF2 appearing in Trichoplax, while AIRE emerging in bony fish. Interestingly, FEZF2 shows a high degree of conservation on the level of functional specificity and positive selection of its proteins. This might indicate that FEZF2 function in vertebrates in part is similar to that of its role in invertebrates. Thus our results might hint toward a role of FEZF2 role in the negative selection of autoreactive immune cells in relation to the innate immune system in invertebrates.

FEZF2 and AIRE have different evolutionary histories. Our results indicate that FEZF2 is more ancient than AIRE as we have found homologs for FEZF2 in Cnidaria and Trichoplax, where the most ancient homolog of AIRE1 exists in fish (figure 2). FEZF2 belongs to the FEZ family and its origin is likely to be a Zinc finger domain-containing protein, C2H2 type (*Metschnikowia aff. Pulcherrima*) (E-value <9e^-32^) (Table 2). FEZF2 still retains these Zinc finger proteins domains (figure 3). Interestingly, zinc finger domains are known to constitute DNA-binding motif in transcription factor TFIIIA however they were recently demonstrated to bind DNA, RNA, protein (28). AIRE1 on the other hand is likely to have evolved from a PHD containing domain gene (figure 3). PHD fingers have been found in various eukaryotic proteins which are responsible for the control of gene transcription and chromatin dynamics confirming the role of AIRE as a transcription factor. Interestingly, PHD fingers can recognize the unmodified and modified histone H3 tail, and some have been found to interact with non-histone proteins(29). This is in agreement with the hypothesis that AIRE1 is capable of regulating transcription factors directly (protein-protein binding) and indirectly through histone modification. Interestingly, although AIRE has emerged in bony fish, it is capable of controlling the expression of more ancient genes such as IGHF2 and SPT1 (figure 2). However, in the case of FEZF2, we have found that it is highly conserved between invertebrates and vertebrates (Table 3). These results imply that FEZF2 regulation of autoreactive genes is conserved in invertebrates hinting at a potential role for FEZF2 regulating simpler mechanisms that might have existed in invertebrates(30). One of the main differences between the genes is their functional motifs (supplementary file 1). Whereas AIRE1 could be binding to nuclear receptors through its motif LXXLL, FEZF2 contains motifs that are involved in autophagy hinting at another role to its known role in the thymus and the brain. It has been shown that FEZF2 plays an important role, in neural development in the brain(14). Autophagy plays a part in sustaining neuronal stem cells and in neuronal plasticity (31). Thus our analysis hints that FEZF2 could be contributing to neural development through the autophagy route. Taken together our results allude that FEZF2 could be contributing to an invertebrates autoimmunity-elimination mechanism through autophagy.

FEZ2 and AIRE1 evolved in Trade-off controlled evolutionary process. Negative selection of auto-reactive T cells is a critical process, which the body needs to ensure its survival. Our study shows a high degree of complementing between FEZF2 and AIRE on several levels (i) evolution history of downstream targets for FEZF2 and AIRE2 (ii) functional ontology of their downstream targets. On the level of evolution history of downstream targets, we found that FEZF2 is far more ancient than AIRE complex (figure 2). However, both of the pathways regulate genes classified as ancient (existing in invertebrates) such as BHMT, F2, ITIH3, KCNJ5, LGALS7, MAOA, MYO15B, PLAG1 and ZP2, in case of FEZF2 and IGF2 and SPT1 in case of AIRE. Interestingly we found downstream targets for both pathways that emerged as late as bony fish, such as ANXA10, APOC2,APOC4, while in case of AIRE, we found C4BP, KRT2 first emerged in fish. On the level of functional ontology of the downstream targets, we found that the two pathways control genes that share the same functional annotation, such as (ZP2 and ZP3). Furthermore, both downstream targets genes share lipid metabolism pathways, as well as genes that play in immune response. Our results indicate that there a high degree of complementary regulation of genes from the same function irrespective of their evolutionary history. We can postulate that the reason behind this complementary approach is to ensure the critical genes that could drive autoimmunity are expressed.

Our results shed light on the negative selection of immune cells in lampreys. We have detected various AIRE and FEZF2 associated genes in lampreys (figure 2). Lampreys evolved with their specific phenotypes of immune cells such as VRLA (homologous α/β T cells), VRLB (homologous to B cells), and VRLC (homologous to γ/δ T cells)(19). Lampreys have a thymoid that expresses VRL (32). This implies that lampreys might have a mechanism for eliminating autoreactive immune cells. Interestingly, we could not locate an AIRE1 homolog in lampreys. However, we found a homolog of FEZF2 in lampreys (XP_032822733.1). We have also detected LTBR, which is known to control FEZF2 expression in lampreys (XP_032823234.1) (13). Similarly, downstream targets, known to be controlled by FEZF2 are also expressed in lampreys such as BHMT, CALCA, COL17A1, CRISP 1, CYP24A1, F2, FABP7, FABP9, GC, ITIH3, KRT10, KCNJ5, LGALS7, LYPD1, Maoa, MUC3, MYO15B, PLAGL1, PLD1,Resp18 and ZP2. Taken together our results indicate that lampreys could possess a negative-selection-like mechanism.

## Conclusion

Our analysis drew a heterogonous picture for the negative selection process. While the AIRE1 pathway has evolved in bony fish, FEZF2 pathways seem to be tightly conserved in invertebrates. The existence of FEZF2 pathways in invertebrates might hint at the conservation of a function in lower organisms. Since T cells are non-existent in invertebrates, our findings indicate that FEZF2 could have been playing a role in a primitive negative selection mechanism of innate immune cells in invertebrates.

## Supporting information

Supplementary file 3-fasta files

## Acknowledgments

We would like to acknowledge the efforts of Macrious Abraham, and Meriam Joachim for their informative discussions.

## Supplementary Materials

Supplementary materials include 1. Motif search results 2. GO results.3.Fasta files

